# Visuomotor skill learning in young adults with Down syndrome

**DOI:** 10.1101/2022.08.22.504780

**Authors:** Laurits Munk Højberg, Jesper Lundbye-Jensen, Jacob Wienecke

**Author notes:** Corresponding author: Laurits Munk Højberg, Department of Nutrition Exercise and Sports, University of Copenhagen, Copenhagen, Nørre Allé 51, 2100 København N, Denmark.

## Abstract

**Background:** Individuals with Down syndrome (DS) have impaired general motor skills compared to typically developed (TD) individuals.

**Aims:** To gain knowledge on how young adults with DS learn and retain new motor skills.

**Methods and Procedures:** A DS-group (mean age = 23.9 ± 3 years, N = 11), and an age- matched TD-group (mean age 22.8 ± 1.8, N= 14) were recruited. The participants practiced a sequence visuomotor accuracy tracking task (VATT). Online and offline effects of practice were assessed in immediate and 7-day retention tests. Participants practiced the task in seven blocks (10.6 minutes).

**Outcomes and Results:** The TD-group performed better than the DS-group in all blocks (all P < 0.001). Both groups improved VATT-performance online from baseline to immediate retention (all P < 0.001). The DS-groups’ performance at 7-day retention was at the same level as the immediate retention tests (ΔDS). An offline decrease in performance was found in the TD-group (ΔTD, P < 0.001). A between-group difference was observed in the offline effect on the sequence task (ΔTD - ΔDS, P = 0.04).

**Conclusions and Implications:** The motor performance of adults with DS is lower compared to their TD peers. However, adults with DS display significant online performance improvement during training, and offline consolidation following motor learning.

**What this paper adds:** Learning new motor skills is fundamental throughout our lifespan. Persons with Down syndrome have other prerequisites for learning new tasks, related to psychological, physiological, and anatomical factors imposed by the syndrome. This study is the first to investigate online and offline learning effects of a single motor skill training session in adults with DS. Our results show generally lower motor performance in DS individuals compared to the typically developed population, but with equal online learning effects. Both groups demonstrate retention, i.e., offline stabilisation but while TD demonstrate negative offline effects, this was not the case for DS. These results should be taken into consideration when planning training of motor and general life skills for adults with DS. This work lays the ground for further investigations of the trajectory of the early learning processes and the mechanisms involved when this target group acquires new skills.

## 1. Introduction

The ability to learn and retain new motor skills is fundamental for everyday functioning, it allows us to perform movements, to meet the demands of our environment, and adapt to changes in it (R. A. Schmidt & Lee, 1999). Individuals born with Down syndrome (DS), are presented with a range of physical and psychological challenges including intellectual disability (Grieco et al., 2015) and impaired general motor skills (Vicari, 2006), which have been attributed to altered anatomical, physiological, and neurological development (Dierssen, 2012). Because of this, persons with DS have a higher need for support from health care and social services throughout their lives (Tsou et al., 2020). Knowledge about the characteristics of the ability to learn, consolidate and retain new motor skills is valuable for the population with DS and professionals working with the population. It can lead to improved planning and structure of motor skill training in the everyday life of individuals with DS (Hall et al., 2011; Jain et al., 2022).

Previous studies have demonstrated that adults with DS can improve their motor performance during practice (Kerr & Blais, 1985; Reilly et al., 2017), and intervention studies have shown that the group can improve their gross motor skills over several training sessions. The latter has been observed as improvements in gait parameters after 10 weeks of Nordic walking (Skiba et al., 2019) and as improved dribbling skills after 16 weeks of soccer practice (Perić et al., 2021). These results confirm the ability of skill learning in people with DS, but the detailed dynamics of online learning (i.e., the change in performance during practice) and the subsequent offline consolidation and retention of motor skills in adults with DS, has to our knowledge not yet been investigated.

Motor skill learning is a complex process, and it is influenced by a range of factors (Dayan & Cohen, 2011), e.g., the organization of task practice (Lage et al., 2015), the type and availability of (augmented) feedback (Oppici et al., 2021) and interindividual differences such as cognitive abilities (Seidler et al., 2013). Both implicit and explicit memory processes are involved in motor skill learning (Hikosaka et al., 2002), and age and baseline skill level seem to influence to which memory system dominate the learning process (Nemeth et al., 2013). Individuals with DS display impairments in explicit memory, and a relatively preserved implicit memory (Vicari et al., 2000), and this distinct memory profile could influence how motor skills are acquired and retained in the population. Motor skill learning is influenced by individual cognitive abilities such as executive functions (M. Schmidt et al., 2017) and spatial working memory (Seidler et al., 2013). Part of the intellectual disability profile of individuals with DS is a deficit in these specific functions (Dierssen, 2012; Lanfranchi et al., 2009), thus the groups’ performance on tests probing these abilities could help explain part of the potential differences in motor skill learning.

On the basis on this knowledge, the present paper addresses the questions: What are the dynamics of online and offline motor skill learning for individuals with DS, and are the dynamics different from TD individuals? Furthermore, can the potential differences in the dynamics of motor skill learning be related to differences in cognitive abilities? To address these questions, a visuomotor accuracy tracking task (VATT), was used. The task involves control of dynamic pinch force in a precision grip, to track visual targets. We evaluated the within-session improvement in performance (online effects) as well as the retention seven days after motor practice (offline effects). We hypothesized that the participants with DS would show improved motor performance during acquisition and display retention seven days later. We expected the typically developed (TD) adults to perform better than the DS-group in the baseline assessment, and that they would display larger online improvements compared to the DS-group, possibly due to the use of explicit processes dominating early skill acquisition.

The task involved five different targets appearing on the screen in a repeated sequence. During motor practice, the sequence was colour-coded, i.e., a specific target colour always appeared in the same position. By implementing two different tests at baseline and retention, one with the sequenced target order, but without colour-coding and one with a random target order, we aimed at investigating to what extent improved motor performance could be related to explicit sequence learning versus improved motor acuity (Shmuelof et al., 2014). The presence of sequence learning was investigated by comparing the performance in the final sequenced practice block (with colours), to the performance in the sequenced, non-colour coded retention block, and by comparing performance in the sequenced and random retention blocks. We expected to observe sequence learning in the TD-group, due to employment of an explicit learning strategy, while we expected this to be absent in the DS-group. To investigate if potential differences in online and offline effects between groups could be related to specific motor or cognitive abilities, we administered a battery of motor and cognitive tests and assessment of fine motor skills.

## 2. Methods

### 2.1 Participants

Twelve adults with Down syndrome (DS-group) and 14 age-matched typically developed adults (TD-group), were recruited. One participant in the DS-group were excluded, since the person were not able to complete the training. Participants’ characteristics is shown in Table 1. The study was approved by The Danish National Committee on Health and Research Ethics (H-19017828) and adhered to the Helsinki Declaration II. All participants gave their informed consent before participation and only participants without guardianship were included.

**Table 1:** Demographic data of the participants

| Variables | DS | TD |
| --- | --- | --- |
| Number of participants | 11 | 14 |
| Sex (f/m) | 3/8 | 6/8 |
| Age (years ± SD) | 23.9 ± 3.0 | 22.8 ± 1.8 |

### 2.2 Experimental design

Each participant visited the lab twice over the course of one week. On day 0 the participants completed the baseline, motor practice and immediate retention blocks. Exactly one week after their first visit, the participants completed 7-day retention tests and additional motor and cognitive tests.

### 2.3 Visuomotor accuracy tracking task

A computer-based dynamic visuomotor accuracy tracking task (VATT) performed with the dominant hand was applied. The task has similarities to tasks used in previously published works from our lab (Beck et al., 2020; Christiansen et al., 2018). The participants were seated approximately 50 cm from a screen at an adjustable table, which was set to ensure a comfortable position of the dominant arm. The tables’ hight was recorded, to ensure replication on the second visit. The task required the participants to control a cursor to track a series of target boxes on the monitor, by accurately applying pinch force to a spring-loaded lever, relaying the force to a load-cell (UU2-K10, Dacell, South Korea) (Figure 1, see also Larsen et al., 2016). Then the force was low-pass filtered (10 Hz), amplified (x100) (AM- 310, Dacell, South Korea), and then fed the PC through a USB-connected data board, sampling at 90 Hz (NI USB-6008, National Instruments, Austin, Texas). An in-house developed Python application ran the VATT. Continuous, online feedback was presented to the participants, as the cursor changed colour from red to blue, when the targets were hit correctly. The targets were presented for 2 s, with 0.2 s between targets. The participants were given augmented feedback, as their time on target in percent were displayed for 2 s after each block. Performance was assessed in two blocks of 59.2 s (27 targets) at baseline, immediately, and seven days after the training session; a block with a random target order and a block with five targets appearing in a repeated sequence. At baseline the random block was performed before the sequence block. The order was reversed at both retention tests. The participants practiced the sequenced task in 7 blocks of 92.2 s each (42 targets), with a 1-minute break between blocks. At baseline and retention, all targets were displayed in red. During practice, the five-target sequence had different colours (augmented information on the sequence), i.e., the same colour always appeared in the same position. Before the first baseline block, the participants completed a sequenced familiarization block with red targets (33 s, 15 targets). The participants were not informed about the differences in block structure. Between the baseline and practice blocks, the participants were seated in rest for 20 minutes. Between practice and immediate retention, the participants had a 1-minute break.

**Figure 1:**
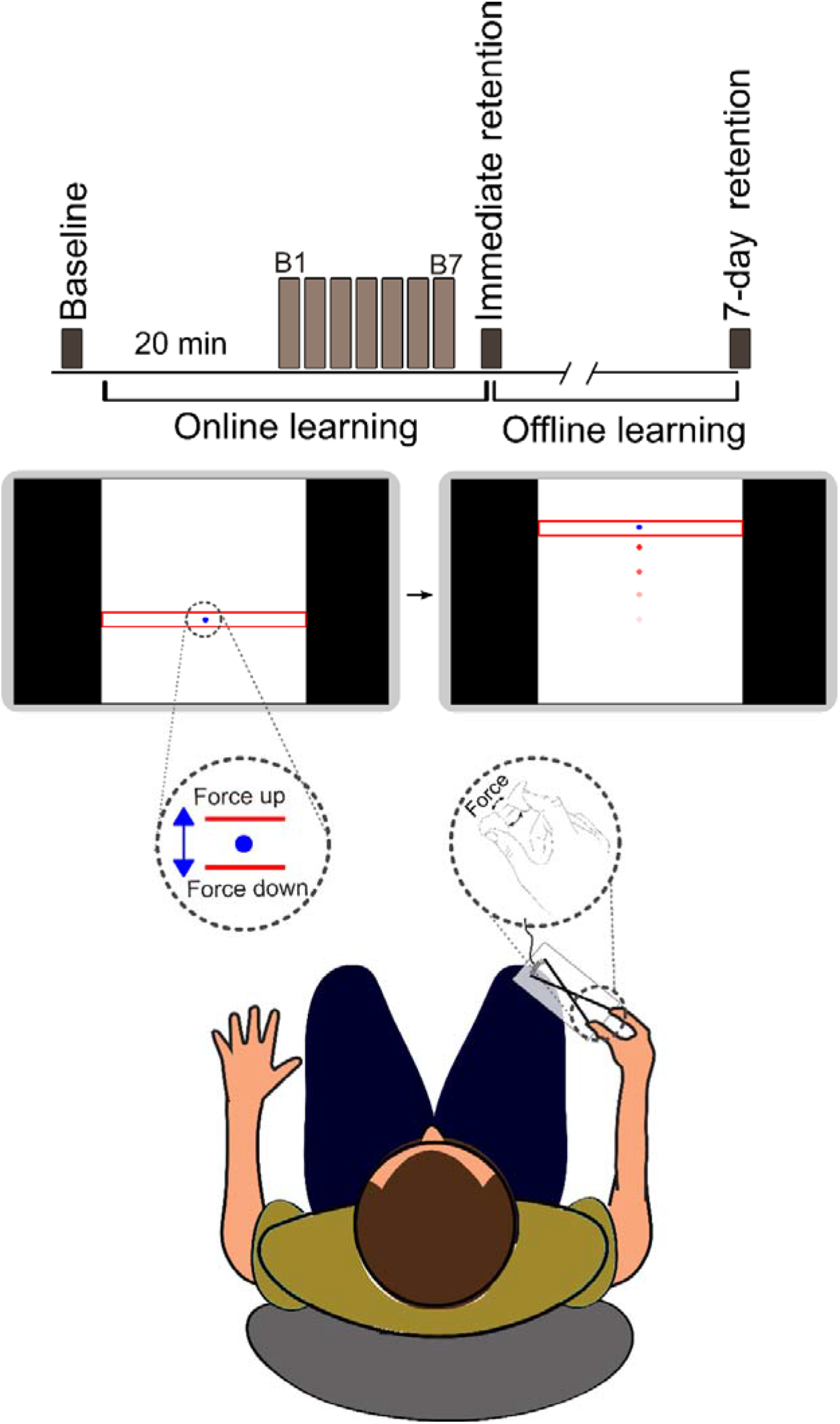
Outline of the VATT and study protocol of the current study. The underarms are resting on an adjustable table, to ensure a comfortable position.

### 2.4 Visuospatial working memory

We applied a computerized forward span corsiblock tapping task, developed by Aeschlimann and co-workers (Aeschlimann et al., 2017). The participants were placed in front of a laptop with a touchscreen with 16 white squares in a 4 by 4 grid. Following a fixation cross, one square at a time changed colours to black in a quasi-randomized pattern, followed by a question mark. Then the participants were instructed to tap the same boxes on the screen in the same order, with their index finger. After an introduction and three practice rounds (two trials with 2-span tasks and one with a 3-span task), the task commenced at 2 spans. Four correct answers out of six trials were required to move on to the next level. If this was not accomplished, the task was terminated.

### 2.5 Basic executive functions

A computerized modified Eriksen flanker task was used to measure basic executive functions (inhibitory control, processing speed, working memory and flexibility, (Eriksen & Eriksen, 1974)). The flanker task applied in this study was originally developed by Adele Diamonds’ research group (Schonert-Reichl et al., 2015). The participants were seated in front of a laptop with an external keyboard. In the task, images of rows of five fish turning left or right was displayed, and the participants were instructed to ‘feed the correct fish as quickly as possible’, either by pressing the left control key with their left index finger or the numpad enter key with their right index finger, determined by the way the fish was facing. The buttons were marked with arrows showing the directions (left ctrl =, numpad enter = ?). The task contained four trial types, determined by the distractors that were either facing the same direction (congruent), the opposite direction (incongruent), upwards (neutral), or without distractors. The task consisted of three sub-tasks: A regular flanker task, a reversed, and a mixed task. In the regular task the fish were blue, and the participants were instructed to feed the middle fish. In the reversed task the fish were pink, and the participants were told to feed the fish on the sides. The mixed task consisted of both blue and pink fish, and the participants were instructed to feed either the middle or the outside fish, depending on the colour of the fish. Before each sub-task, the participants completed practice rounds. In the regular and reversed tasks, the practice round consisted of 4 stimuli. Another practice round was required if 2 or less trials were correct. The mixed practice round consisted of 8 stimuli, and another round were performed if a participant got 5 or less correct answers. A maximum of three practice rounds were allowed for each sub-task. The test was terminated if a participant failed the third practice round before any the sub-tasks. The regular and reversed tasks consisted of 17 stimuli each (3 congruent, 10 incongruent, 2 neutral and 2 without distractors), and the mixed task consisted of 45 trials (12 congruent, 16 incongruent, 9 neutral and 8 without distractors). The average reaction times (RT) on all trials and the RTs on the congruent and incongruent trials were extracted as outcomes, as well as the accuracy rates on the congruent and incongruent trials (correct answers/total number of trials). The flanker effects were calculated as: incongruent – congruent, both in the RT and accuracy measures.

### 2.6 Test of fine motor skills

The Purdue Pegboard test (Lafayette Instrument Company, Lafayette, Indiana, USA) were performed to measure the participants’ manual dexterity (Tiffin & Asher, 1948). The test consisted of four sub-tasks: 1) The participants placed as many pins in the board as possible in 30 seconds with their dominant hand. 2) Similar to 1), but with the non-dominant hand. 3) The participants worked bimanually and had to place as many pairs of pins as possible in 30 seconds. 4) The assembly task, where the subjects were instructed to build as many assemblies as possible using three different parts in 60 seconds. Before each task, the experimenter instructed and demonstrated the task, and the participants practiced the task. Outcomes were the Sum of Scores (the sum of correctly placed pins in sub-task 1), 2), and 3)) and the Assembly score (the number of correctly placed parts in subtask 4)).

### 2.7 Data analysis and statistics

All analyses were performed in R statistics (*R Statistics 4.0.0*, 2020). A linear mixed effects model was fitted to the data (“lme4” R-package (Bates et al., 2015)), to investigate the effect of time (i.e., the performance on the 13 blocks), group and type of block (sequence or random) on motor performance. Time on target in seconds, were translated to a score between 0 and 100 points, with each point being equal to 0.02 s. Group, time, and block type were included as fixed effects, together with sex and corsiblock span performance (the level at which the participants failed the test). Sex and corsiblock performance were included to control for the variation these factors could introduce to the data. Since the participants were likely to display differences in baseline performance and our experiment is a repeated measures design, random intercepts were fitted for each participant.

(1): Time on target ∼ Group x Block x Type + Sex + Corsiblock span number + (1|participant)

The equation above is presented in R-terminology where “(1|participant)” represents the individual intercepts and the x’es indicates the interactions between the fixed effects of Group, Block and Type. If significant effects of the interactions between Time on target and Group and Block (time), were observed, post hoc analyses of within- and between-group differences were performed. The post hoc analyses investigated differences in online and offline leaning, and differences in performance between the random and sequence blocks, to investigate sequence learning. P-values from the post-hoc analyses were corrected for multiple comparisons with the single step method. Paired T-tests were used to investigate differences between the groups in the motor and cognitive tests. Unpaired T-tests were used to determine the presence of flanker effects within the groups. Alpha level was set at P < 0.05, for all analyses. Results from the linear mixed effects model are presented as model estimated means ± standard error (SE). The results of the motor and cognitive test are presented as mean ± SE. Correlation matrixes were computed for each motor and cognitive tests and VATT performance on the sequenced task at baseline and online and offline effects, and for the absolute motor performance level on day 1 and offline effects.

## 3. Results

### 3.1 Fine motor skills and cognitive test performance

Paired T-tests showed that the TD-group performed better in the pegboard task and the corsiblock tapping task compared to the DS group (all P-values < 0.001, Table 2). The results from the flanker tasks showed that the TD-group had lower reaction times (RT) on the three sub-tasks, both in the average RT for all trials and on the congruent and incongruent trials (table 2): Regular flanker (all P-values < 0.001), reversed flanker (all P-values < 0.01) and mixed flanker (all P-values < 0.05). The TD-group was more accurate on the incongruent trials in all three tasks compared to the DS group (all P-values < 0.01), but no differences were observed on the congruent trials. No flanker effects were observed for RT in either of the groups in the regular flanker (DS-group: P = 0.923, TD-group: P = 0.192). The DS-group displayed a flanker effect in accuracy in the regular flanker task (P = 0.006), while the TD-group did not (P = 0.104), this was seen as a significant difference between the groups in the paired T-test (P < 0.05). In the reversed flanker task, the DS-group presented a flanker effect on RT (P = 0.025), while the TD-group did not (P = 0.083). The paired t-test revealed a significant difference in the flanker effect on RT between the groups (P = 0.019). With regards to accuracy, both groups displayed flanker effects (DS-group: P = 0.045, TD-group: P = 0.040). In the mixed task, both groups presented flanker effects on RT (DS-group: P = 0.016, TD-group: P = 0.001), and accuracy (DS-group: P < 0.001, TD-group: P = 0.012). The paired T-tests between the groups showed no difference between the groups in the flanker effect in RT (P = 0.105), and a significant difference between the groups in the flanker effect on accuracy (P < 0.001).

**Table 2:** Outcomes of the motor and cognitive tests

| Variables | DS |  | TD |  | Sig. level |
| --- | --- | --- | --- | --- | --- |
| Pegboard |  |  |  |  |  |
| Sum of scores | 18.5 ± | 1.7 | 39.2 ± | 1.0 | *** |
| Assembly | 13.1 ± | 1.3 | 43.7 ± | 1.4 | *** |
| Flanker test |  |  |  |  |  |
| Regular RT all (ms) | 1400 ± | 160 | 464 ± | 17 | *** |
| Regular congruent RT (ms) | 1376 ± | 176 | 469 ± | 22 | *** |
| Regular incongruent RT (ms) | 1406 ± | 160 | 454 ± | 16 | *** |
| Regular congruent acc. | 0.83 ± | 0.07 | 1 ± | 0 |  |
| Regular incongruent acc. | 0.45 ± | 0.11 | 0.96 ± | 0 | ** |
| Regular flanker effect RT (ms) | 26 ± | 123 | -16 ± | 13 |  |
| Regular flanker effect acc. | -0.38 ± | 0.11 | 0.04 ± | 0 | * |
| Reversed RT all (ms) | 1299 ± | 123 | 439 ± | 132 | *** |
| Reversed congruent RT (ms) | 1101 ± | 1587 | 471 ± | 26 | ** |
| Reversed incongruent RT (ms) | 1338 ± | 126 | 446 ± | 19 | *** |
| Reversed congruent acc. | 0.93 ± | 0.06 | 1 ± | 0 |  |
| Reversed incongruent acc. | 0.82 ± | 0.04 | 0.98 ± | 0 | ** |
| Reversed, flanker effect RT (ms) | 290 ± | 115 | -25 ± | 9 | * |
| Reversed, flanker effect, acc. | -0.11 ± | 0.06 | -0.02 ± | 0 |  |
| Mixed, RT all (ms) | 1085 ± | 77 | 730 ± | 35 | ** |
| Mixed congruent RT (ms) | 934 ± | 577 | 683 ± | 35 | ** |
| Mixed incongruent RT (ms) | 1198 ± | 125 | 799 ± | 38 | * |
| Mixed, congruent acc. | 0.92 ± | 0.05 | 0.97 ± | 0 |  |
| Mixed, incongruent acc. | 0.47 ± | 0.06 | 0.92 ± | 0 | *** |
| Mixed flanker effect, RT (ms) | 268 ± | 104 | 117 ± | 27 |  |
| Mixed flanker effect, acc. | -0.45 ± | 0.04 | -0.05 ± | 0 | *** |
| Corsiblock tapping task |  |  |  |  |  |
| Span number | 2.8 ± | 0.4 | 6.4 ± | 0.4 | *** |
| Correct answers | 6.9 ± | 1.8 | 26.5 ± | 1.7 | *** |

Sum of scores = The sum of pins placed with the right, left and both hands in three separate trails. Assembly score = The number of correctly placed parts in the assembly task. RT = Reaction time. Acc. = Accuracy ratio, calculated as number of correctly answered trials divided by the number of total trials. Flanker effect RT = Reaction time in the congruent trials subtracted from the reaction time in incongruent trials. Flanker effect acc. = Accuracy ratio in the congruent trials subtracted from the accuracy ratio in the incongruent trials. Corsiblock Span number = The number of spans at which the participants failed the task. Correct answers = The number of correctly performed trials. Data is presented as means ± SE. Stars indicate the P-values of the paired T-tests between the groups. Significance codes: "\*" denotes $p < 0.05$ , "\*\*\*" denotes $p < 0.01$ , "\*\*\*\*" denotes $p < 0.001$ .

### 3.2 Motor performance in the VATT

Main effects of both Block (F = 29.6, P < 0.001) and Group were observed (F = 30.1, P < 0.001), so post hoc analyses were performed. The post hoc analyses revealed that the TD-group performed ∼30 points better than the DS group at all timepoints (all P < 0.001), equivalent of 0.6 seconds time on target on each target (Figure 2). A significant effect of sex was found (F = 4.6, P = 0.04) in the VATT, while no significant effect of corsiblock performance were observed (P = 0.88).

**Figure 2:**
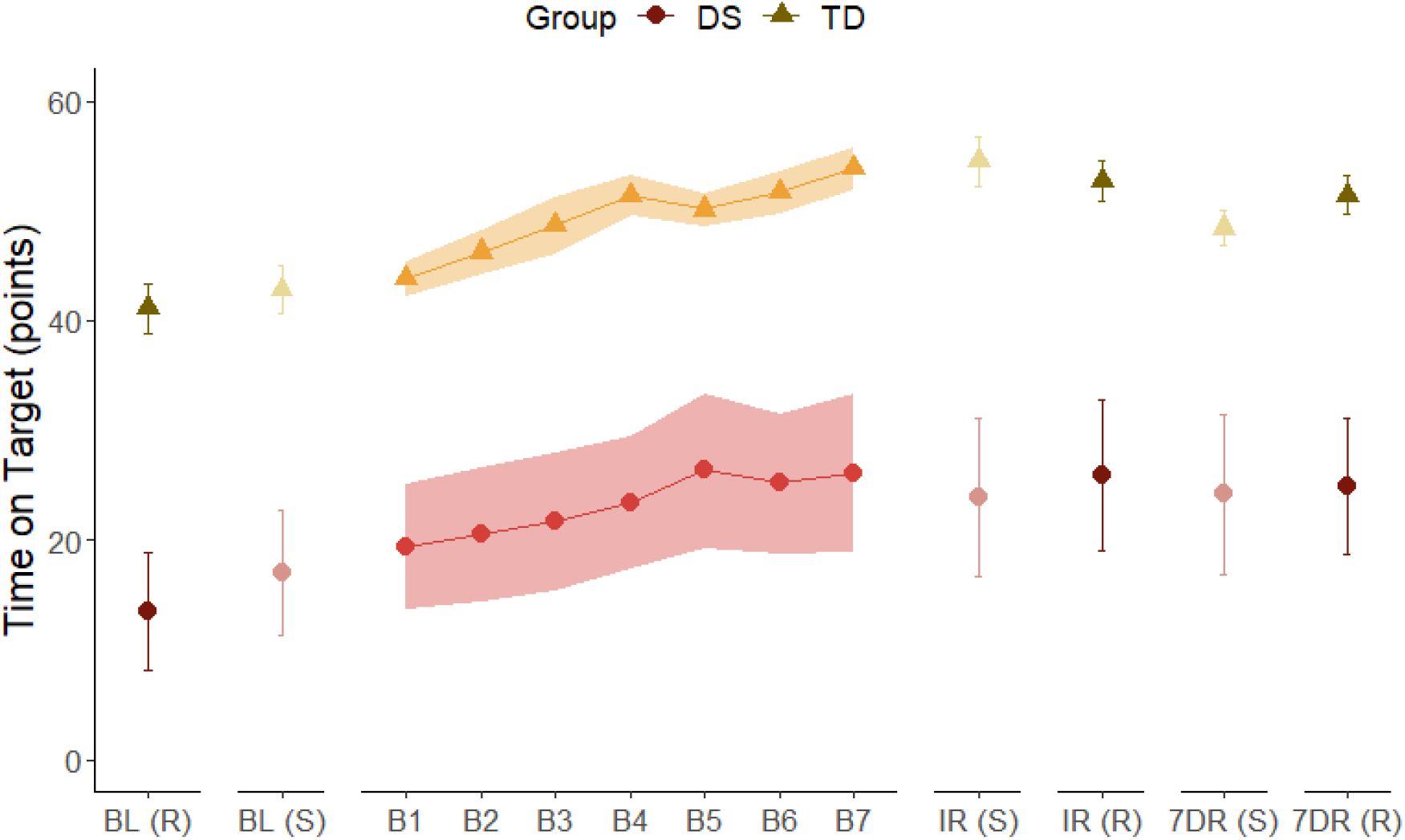
Motor Performance at baseline, during acquisition and at retention tests. Motor performance as percentage points (pp) time on target on all blocks. Black and grey colours indicate random and sequence blocks at baseline, immediate and 7-day retention, respectively. Coloured blocks indicate acquisition blocks. Error bars and ribbons represent error as confidence intervals. Abbreviations: BL = Baseline; IR = Immediate retention; 7DR = 7-day retention, (S) = Sequence; (R) = Random.

### 3.3 Online effects

Changes in visuomotor performance are presented as changes in time on target score. The learning curves for both groups are depicted in Figure 2. The improvements in motor performance within session (online) and between sessions (offline) are shown on Figure 3. Both the groups improved their online motor performance in the sequence task from baseline to immediate retention, (immediate retention – baseline, Δonline) (Δonline DS = 8.4 ± 1.7, P <0.001; Δonline TD = 11.9 ± 1.5, P < 0.001, Figure 3 A and B). No difference in online learning was observed between the groups (Δonline TD – Δonline DS).

**Figure 3:**
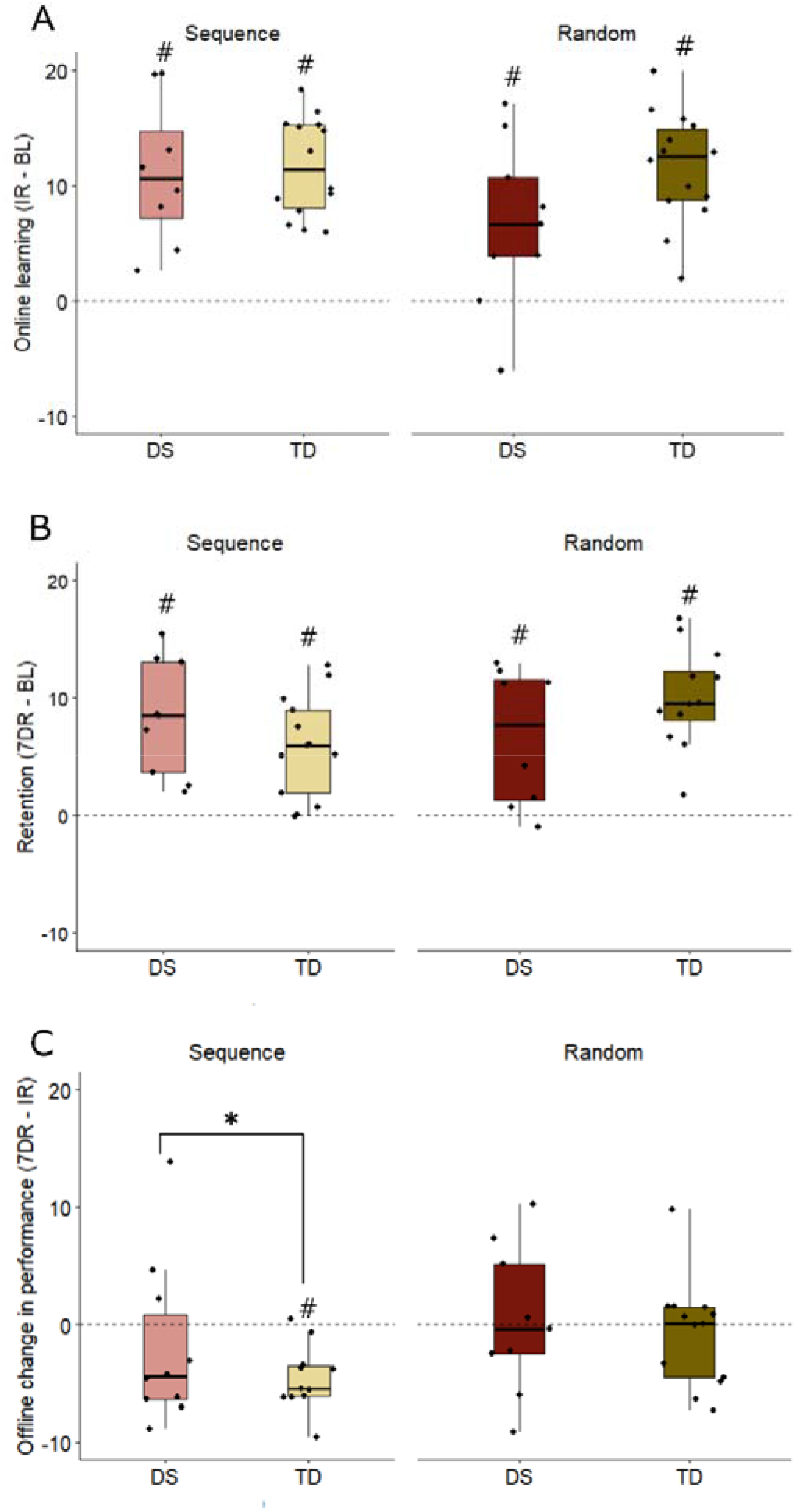
Online and offline effects. Box plots with individual values displaying changes in performance on both the sequenced and random tasks. (A) online performance change from baseline to immediate retention, (B) retention of the VATT, measured as performance changes from baseline to 7-day retention, and (C) offline change in performance from immediate to 7-day retention. The dotted lines at 0 on the y-axes is equal to the groups’ motor performance on the baseline blocks (A and B) and immediate retention blocks (C). BL = Baseline, IR = Immediate retention, 7DR= 7-day retention. # = Significant within-group difference. * = Significant between-group difference in the relative changes.

### 3.4 Retention and Offline effects

Both groups showed retention of the task, as their performances on the 7-day retention tests were significantly better than their baseline performances on both the sequence (DS: 7.9 ± 1.7, P < 0.001, TD: 5.8 ± 1.5, P < 0.001, Figure 3B) and random tasks (DS: 11.3 ± 6.7, P < 0.001, TD: 10.9 ± 1.5, P < 0.001, Figure 3B). No difference between the groups was observed in retention. No difference in motor performance was observed in the DS-group from immediate to 7-day retention in the sequence task (7-day retention – immediate retention, Δoffline) (Δoffline DS: -0.5 ± 1.7, P = 0.995, Figure 3C). The TD-group had a significant offline decrease in performance in the sequence task (Δoffline TD: -6.1 ± 1.5, P < 0.001, Figure 3C). A between-group difference was observed in the offline effects in performance in the sequence task from immediate to 7-day retention (Δoffline TD – Δoffline DS: -5.6 ± 2.2 P = 0.04, Figure 3C).

### 3.5 Implicit and explicit learning

The amount of explicit sequence learning vs. implicit learning of the task, was investigated within groups by analysing performance differences between the sequence and colour-coded practice block 7, and both the sequenced non-colour-coded and the random immediate retention blocks. This was also done between the retention blocks at day 7. No significant differences in performance within or between groups were observed in these analyses.

### 3.6 Correlations between online performance and offline effects

A correlation matrix between motor performance on the immediate retention blocks and the performance changes from immediate and 7-day retention, showed significant negative correlations between the offline change in performance on the sequence task and absolute motor performance in both the random (R = -0.48, P = 0.020) and the sequence immediate retention block (R = -0.57, P = 0.005).

### 3.7 Correlations between motor performance and motor and cognitive tests

The correlation matrix for the flanker task showed a significant negative correlation between baseline VATT performance and reaction time on the congruent trials in the regular flanker task (P = 0.005; Sequence: R = -0.75, P < 0.001), and a significant positive correlation between accuracy on the incongruent trials and baseline VATT performance (R = 0.68, P < 0.001). In the flanker effects, we observed significant positive correlations between the baseline VATT performance and the flanker effects on accuracy on the regular (R = 0.46, P =0.031) and mixed flanker task (R = 0.75, P < 0.001). We did not observe any correlations between baseline VATT performance and the flanker effects on reaction time. In the Pegboard task, significant positive correlations were observed between baseline VATT performance and both Sum of Scores (R = 0.78, P < 0.001) and Assembly (R = 0.85, P < 0.001). Positive significant correlations were observed between baseline VATT performance and corsiblock performance (span number: R = 0.75, P < 0.001, correctly answered trials: R= 0.78, P < 0.001). The correlation matrixes also included the online and offline effects. No correlations were observed between online or offline effects and outcomes from the flanker, Pegboard or corsiblock tasks (P-values > 0.05).

## 4. Discussion

This study is the first to investigate motor skill learning in a task that requires continuous dynamic control of pinch force in adults with DS, and the first to investigate both online and offline effects of motor practice.

### 4.1 Online effects and sequence learning

The participants in both the DS- and TD-group demonstrated positive online effects of motor practice i.e., improved their performance from baseline to immediate retention tests. This finding is in line with previous studies, which have shown that individuals with DS can improve their motor performance during practice (de Mello Monteiro et al., 2017; Kerr & Blais, 1985, 1987). Kerr and Blais used a pursuit tracking task performed with a steering wheel, which required the participants to accurately move the wheel to a given target location to investigate probability learning in youths with DS (Kerr & Blais, 1985, 1987). The pursuit tracking task somewhat resembles the visuomotor tracking element of the VATT in the present study. The researchers observed online performance improvements as a ∼23% decrease in total trial time in 8 blocks and 800 trials (from ∼2300 ms to ∼2080 ms) (Kerr & Blais, 1985), while the present study shows a 40% increase in time on target in 7 blocks with 294 trials (from 17.03 to 23.88, Figure 2). Direct comparisons in the magnitude of the improvements should be made with care, as the tasks are innately different. The aim of the pursuit tracking task was to investigate probability learning in persons with DS, not how the group acquired the motor skill of steering the wheel in a precise manner, while the aim with the VATT in the present study was to assess changes in the ability to control pinch force. Despite differences in baseline motor performance, the DS- and TD-group showed equal improvements in online motor performance. No previous study has reported similar findings. In some previous studies, the age-matched control groups did not improve their performance in the tasks; they performed at maximum level at the baseline test leaving no further room for improvement i.e., a ceiling effect (de Mello Monteiro et al., 2017; Kerr & Blais, 1985, 1987). This discrepancy in online effects between the present and earlier studies could be explained by differences in task designs, including difficulty levels. For instance, the study by Monteiro and colleagues applied a prediction reaction time model, which seem to leave very little room for improvements in the TD-group. The researchers also divided the DS-group into a high and low performing group based on their baseline scores. They observed that only the low performing group improved their scores. The requirements for precise pinch force control in the visuomotor tracking task of the present study, was chosen deliberately, as the aim was to investigate potential differences in motor skill learning between the groups, and that requires a task which challenges participants in both groups.

In addition to motor acuity, the task of the present study involves a prediction and reaction element as well; the faster the participants move the cursor to a target when it appears, the higher a score they can obtain. Thus, it could be speculated that part of the differences between the groups could be attributed to differences in RT and potentially to prediction in the sequence-based task. To our knowledge no previous study has applied such a model in individuals with DS. The flanker task used in the present study includes an RT-element. Here, we find significantly higher RTs on all three tasks in the DS-group compared to the TD-group, ranging from ∼950 ms in the regular task to ∼300 ms in the mixed task, which is in line with previous findings (Davis et al., 1991). This difference in reaction time could partly explain the difference in VATT-performance of the DS-group and the TD-group, however, further experiments and analyses are necessary to conclude on the role of differences in motor reaction time in the VATT.

The protocol of the present study was chosen based on pilot experiments, which indicated that protocols similar to previous studies (e.g., 6 blocks of 3 minutes) was exhausting for the participants with DS to complete (Beck et al., 2020; Thomas et al., 2016). We expected to observe sequence learning after exposure to the colored sequence during the seven practice blocks in the TD-group, as this has been observed previously (Krakauer et al., 2019; Shea et al., 2006). However, no differences in performance between Block 7 (sequenced and colour-enhanced) and the sequence block at immediate retention (sequenced, un-coloured), nor between the sequence and random block at immediate retention was observed in the TD-group. We did not observe sequence learning within the DS-group as expected. It could be the reduced amount of total practice time and less exposure to the sequence that is the cause of the absence of sequence learning in the TD-group. If the participants had been told to be aware of a sequence in the task, it is possible that sequence learning would have been observed.

The results for online effects demonstrate that individuals with DS improve performance with motor practice to the same extent as TD individuals. Since initial skill acquisition is influenced by involvement of cognitive processes and explicit aspects of learning (Krakauer et al., 2019) we could have expected higher performance gains in the TD-group since this group is characterized by higher performance in the cognitive tests. This was however not the case. While the similar online effects between the two groups may be influenced by the higher baseline performance in the TD-group, it can nevertheless be concluded that DS-group demonstrate skill acquisition.

### 4.2 Offline effects, consolidation, and retention

To investigate offline effects and retention on motor learning, we employed delayed retention tests seven days after acquisition of the VATT. The present study is, to the authors knowledge, the first to apply such a design in adults with DS. Earlier studies have demonstrated that persons with DS improve both their gross (Almeida et al., 1994; Perić et al., 2021; Skiba et al., 2019) and fine motor performance (Latash et al., 2002) with several practice sessions practice. The previous studies did not include performance measures between sessions; thus, it is not possible to gauge the offline effects or retention of the motor practice between single sessions. The results of the present study show that adults with DS exhibit retention on the VATT, as significant increases in motor performance at 7-day retention compared to the baseline performance. The DS-group exhibited the same level of retention as the TD-group. A difference between the groups in the offline changes in motor performance on the sequence task was observed (Figure 3B). The DS-group maintained their performance from immediate to 7-day retention, while performance decreased in the TD-group. This difference between the groups could be explained by the difference in absolute performance, which was significant on all blocks throughout the study (∼0.6 s time on target per target). Indeed, we observed negative correlations between absolute motor performance on both the retention blocks on day 1 and the offline change in motor performance in the sequence task, indicating that a high absolute performance on day 1, correlated with a larger decrease in performance from immediate to 7-day retention. The TD-group might have reached a performance close to the ceiling of the task, thus making it less likely for the group to achieve the same level of performance seven days after the acquisition. Reis and colleagues observed either a performance maintenance or decrease on a continuous pinch task, suggesting that this offline pattern might be the default on continuous pinch tasks, contrary to serial tapping tasks, where offline increases in performance have been observed (Reis et al., 2009; Siengsukon & Al-Sharman, 2011).

Additionally, the functional task difficulty, i.e., the difference in task difficulty imposed by differences in skill level of the participants is important to consider (Guadagnoli & Lee, 2004). We observe a significant difference in absolute skill level throughout the experiment, with the TD-group performing at a higher level. The difficulty level of a task during acquisition is related to performance level at retention in a in inverted U-shaped manner, where both a too easy and too hard task is detrimental to retention of the task (Akizuki & Ohashi, 2015). It could be speculated that the task was closer an optimal challenge point for the DS-group, while it was too easy for the TD-group, and thus the negative offline effect was observed. To address the issue of functional difficulty level, future studies should include tasks, that are adjusted to ensure the same absolute performance level at baseline, and a control group that is matched to the same absolute online performance level as the DS-group. The differences in offline effects between the groups could point to differences in how the explicit and implicit memory systems are engaged during online motor learning (Robertson et al., 2004). Previous research has shown that individuals with DS have deficits in explicit memory and implicit memory abilities comparable to typically developed peers (Vicari et al., 2000). It could be speculated that the DS-group employs a predominantly implicit learning strategy during skill acquisition, and that could be part of the reason for the maintained motor performance observed in the DS-group. The TD-groups’ decrease in visuomotor performance at 7-day retention could indicate the engagement of explicit, cognitive processes that is prone to interference due to the presentation of competing knowledge during the offline period (Fletcher et al., 2005). However, we did not find indications of explicit sequence learning on day 1, which contradicts the notion that it is a loss of explicit knowledge of the sequence that is the cause of the performance decrease in the TD-group.

### 4.3 Variability in performance and motor and cognitive tests

Several factors could contribute to the difference in general motor performance level between the groups in the present study. Recent studies have demonstrated increased accuracy and lower variability in the pinch force task with increased age during neurotypical development (Beck, Spedden, & Lundbye-Jensen, 2021; Beck, Spedden, Dietz, et al., 2021). Studies investigating the ability to modulate or hold a specific force output have shown greater variability and a generally lower force output in individuals with DS (Heffernan et al., 2009; Rao et al., 2017). In the present study, we observe greater interindividual variability in the DS-group, than in the TD-group, evident as larger confidence intervals in Time on Target score (Figure 2). In addition, children and adolescents with DS have a different hand motor control development with specific grasping characteristics; they generally grasp objects with fewer fingers and the fingers not used for the actual grasping are often extended (Jover et al., 2010). These previous findings taken together with the results from the motor and cognitive tests of the present study, could explain the difference in visuomotor performance observed between the groups: Higher reaction times along with difficulties in controlling the pinch force will reduce time on target in the VATT. We observed significant correlations between baseline VATT performance, and performance in the pegboard, corsiblock, and flanker tasks. This indicates a relation between performance on the VATT, and the skills measured by these tests, while they also are an expression of the general cognitive and motor challenges related to DS.

### 4.4 Limitations

An investigation of the presence of explicit knowledge about the motor sequence, e.g., as a questionnaire would have been advantageous, as it would have provided an indication of learning strategies. It is important to note that the observed difference between the groups in offline effects, needs to be investigated further. Indeed, the maintenance of motor performance in the DS-group seems to be driven by a large offline improvement in one participant and small improvements in three, while the remaining seven participants reduce their performance roughly by the same amount as the TD-group (Figure 3C). Despite this, our data analysis demonstrates a statistically significant difference between the groups. Lastly, it would have been of interest to have information on the level of severity of the intellectual disability of the DS-group, as this could to influence the performance of motor tasks (Gimenez et al., 2017).

## 5. Conclusion

Motor performance in the DS-group was lower compared to the TD-group throughout the experiment. The groups displayed similar improvements in online motor performance. The inclusion of a delayed retention test in a motor learning scenario for young adults with DS is novel and allowed an investigation of how a new motor skill is retained in this population. Both groups demonstrated significant retention seven days after motor practice. While the TD-group displayed a decrease in performance at 7-day retention in the sequence task from the groups’ immediate retention performance level, motor performance was maintained in the DS-group and a difference between the groups was observed. These findings demonstrate that individuals with DS can indeed acquire and retain motor skills with practice. The cause for the difference in offline effects between the groups is unclear, but we suggest it is the result of the difference in absolute performance and functional difficulty level. The results did not indicate any sequence learning in the groups, as no differences in motor performance between the last sequence-coloured acquisition block and the sequenced non-coloured retention block or the random retention block were observed.

